## Supplementary information for "SWITCH 1/DYAD is a novel WINGS APART-LIKE antagonist that maintains sister chromatid cohesion in meiosis"

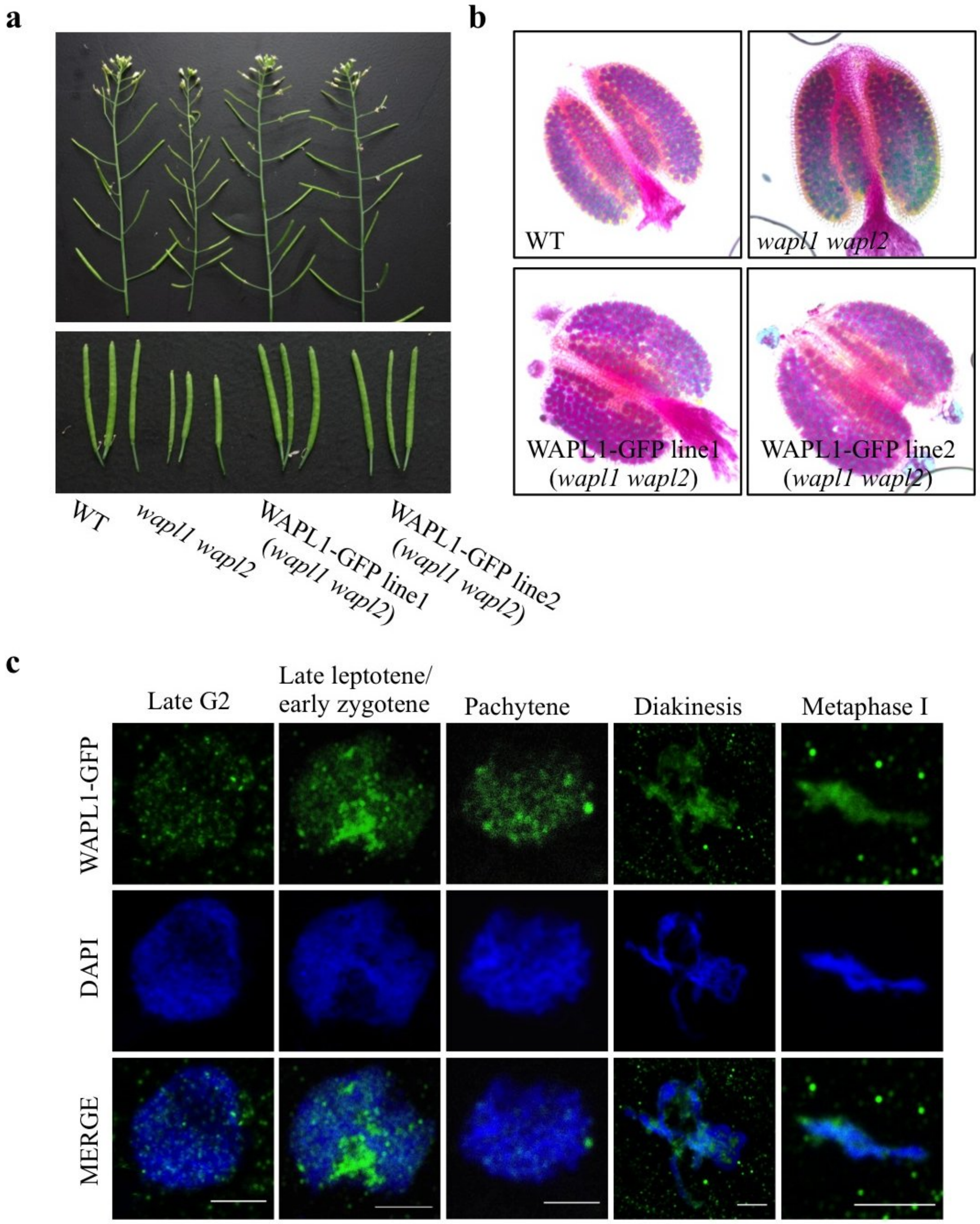


**Supplementary Figure 1**

**WAPL1-GFP is functional and accumulates on chromatin from late leptotene/early zygotene till metaphase I.** (a) Main branches (upper panel) and siliques (lower panel) of the wildtype (WT), *wapl1 wapl2*, and two lines expressing WAPL1-GFP in a *wapl1 wapl2* mutant background. (b) Peterson staining of anthers in the wildtype (WT), *wapl1 wapl2* mutants, and two WAPL1-GFP lines. Blue staining indicates dead pollen. (c) Immunolocalization of WAPL1-GFP during meiosis I of male meiocytes. Anti-GFP antibody was used for detecting WAPL1-GFP. Bar: 5 μm.

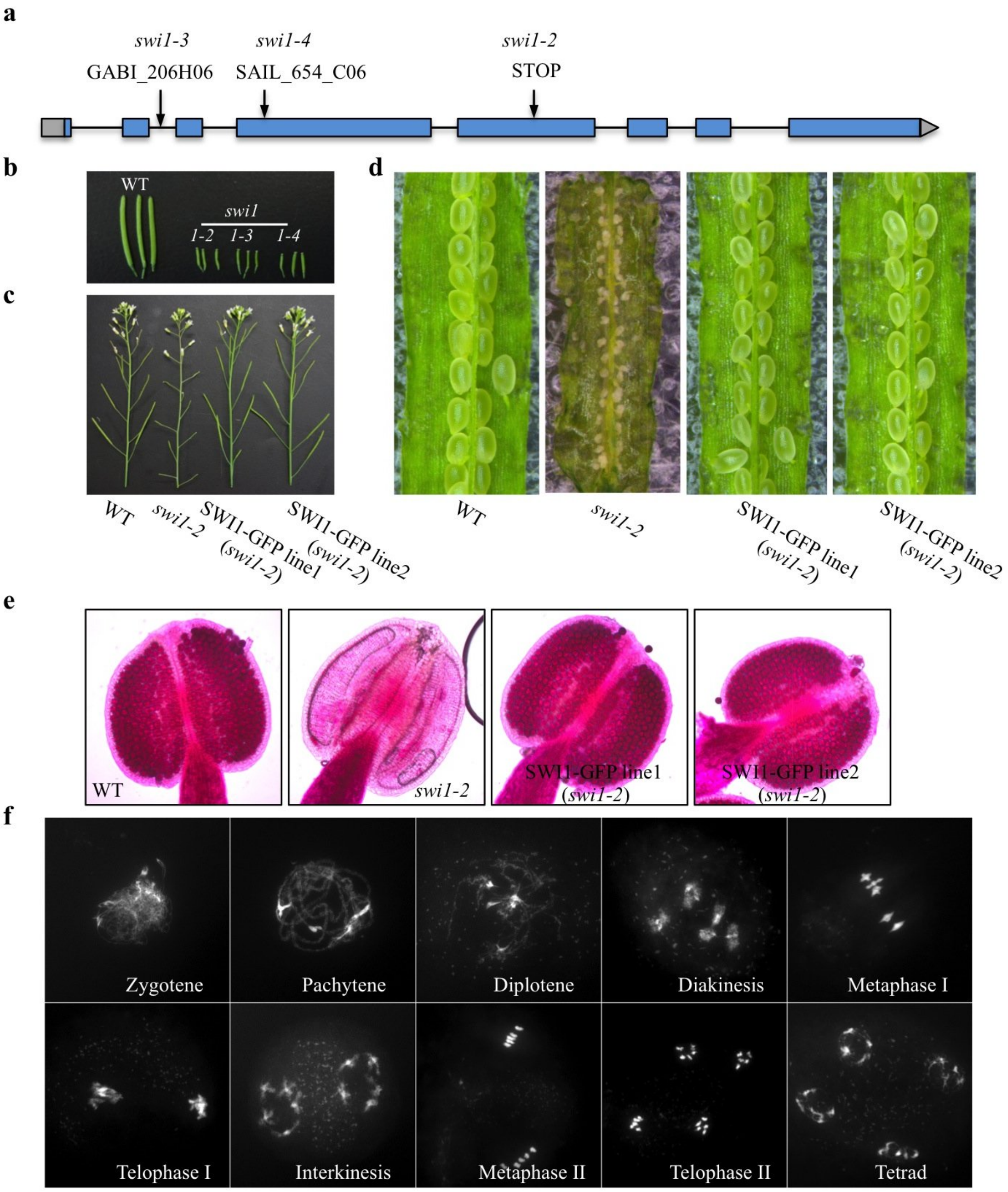


**Supplementary Figure 2**

**SWI1-GFP fully complements the meiotic defects in *swi1* mutants.** (a) Scheme of the genomic region of SWI1. Arrows denote the position of T-DNA insertions (*swi1-3*, *swi1-4*) and of a premature stop codon (*swi1-2*). (b) Siliques of the wildtype (WT) and different *swi1* mutant alleles, which are completely sterile. (c) The main branches of the wildtype, *swi1-2* and two *SWI1-GFP* lines. (d) Seed sets in siliques of the wildtype, two *SWI1-GFP* lines and the *swi1-2* mutant. (e) Peterson staining of pollen for the wildtype, *swi1-2* and *SWI1-GFP* lines. No pollen was found in the *swi1-2* mutants. Blue staining indicates dead pollen. (f) Chromosome spread analysis of male meiocytes in *SWI1-GFP* line #2 (*swi1-2*) shows a wild-type meiotic program.

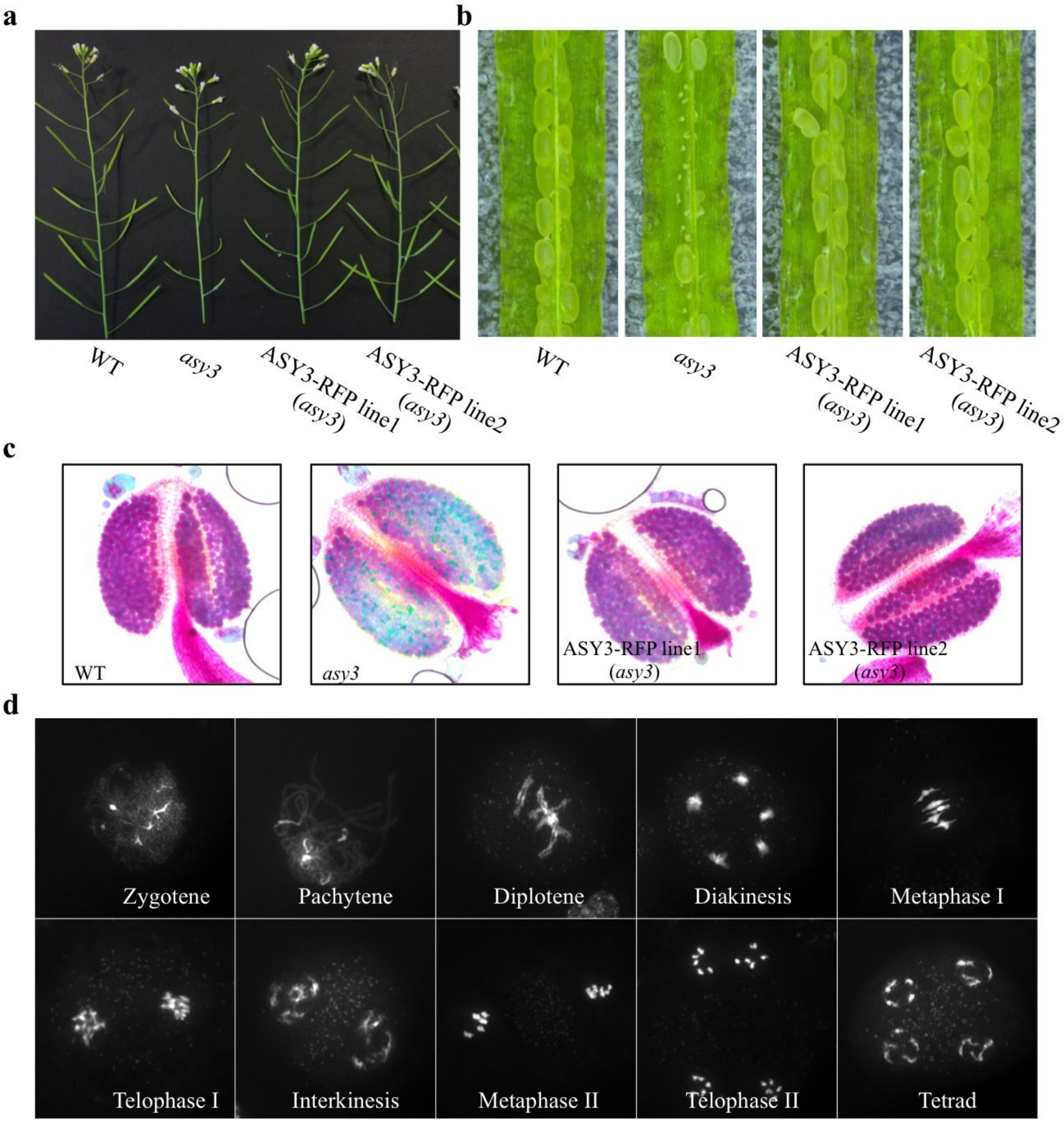


**Supplementary Figure 3**

**ASY3-RFP fully complements the meiotic defects in *asy3* mutants.** (a) The main branches of the wildtype (WT) and two *ASY3-RFP* lines. (b) Seed sets in siliques of the wildtype (WT) and two *ASY3-RFP* complementary lines. (c) Peterson staining of pollens for the wildtype (WT), *asy3* and *ASY3-RFP* lines. (d) Chromosome spread analysis of male meiocytes in *ASY3-RFP* line #1 (*asy3*) reveals a normal meiotic program.

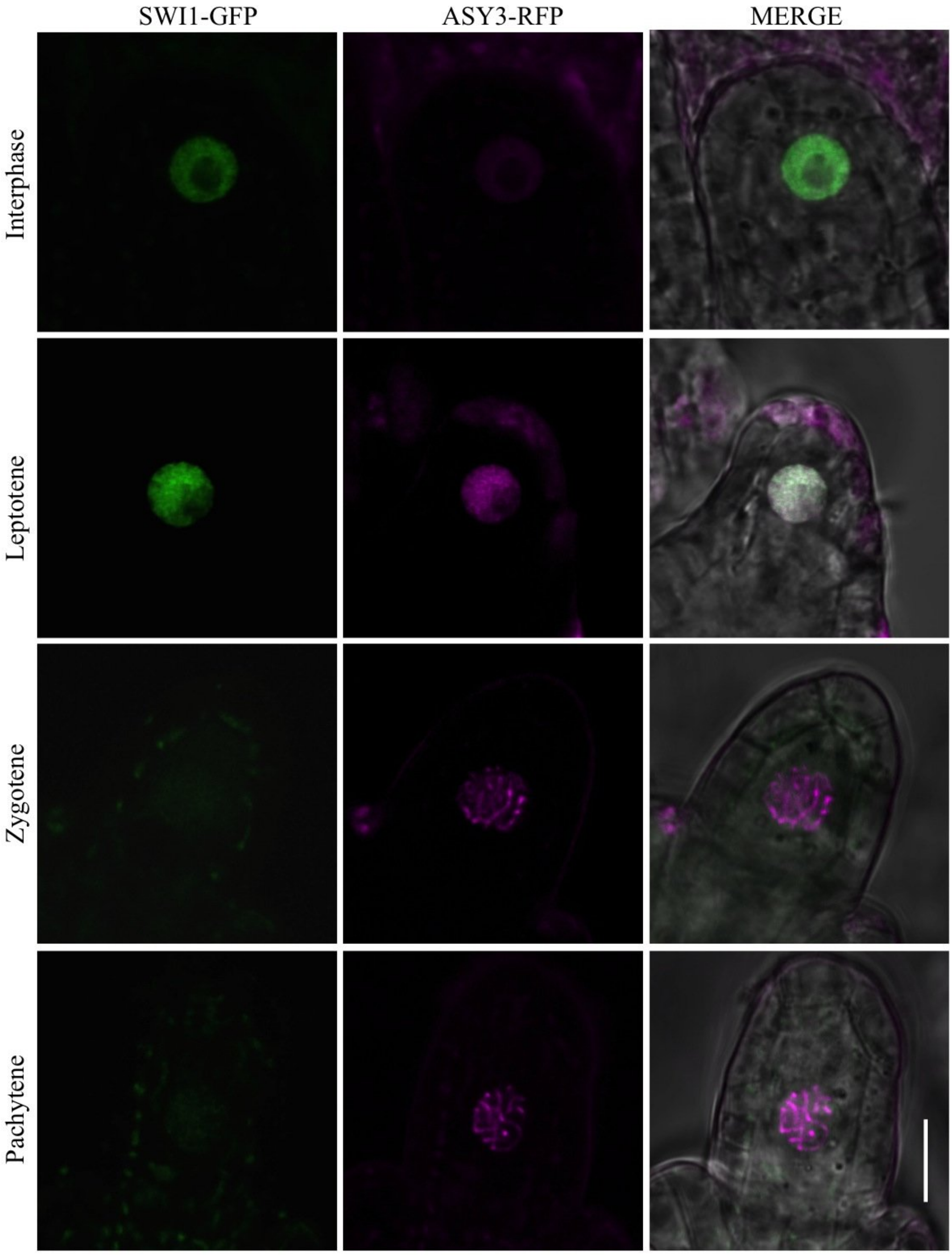


**Supplementary Figure 4**

**Co-localization of SWI1 with ASY3 in female meiocytes.** Co-localization analysis of SWI1-GFP (green) with ASY3-RFP (red) during interphase and prophase I in female meiocytes by using confocal laser scanning microscopy. Bar: 10 µm.

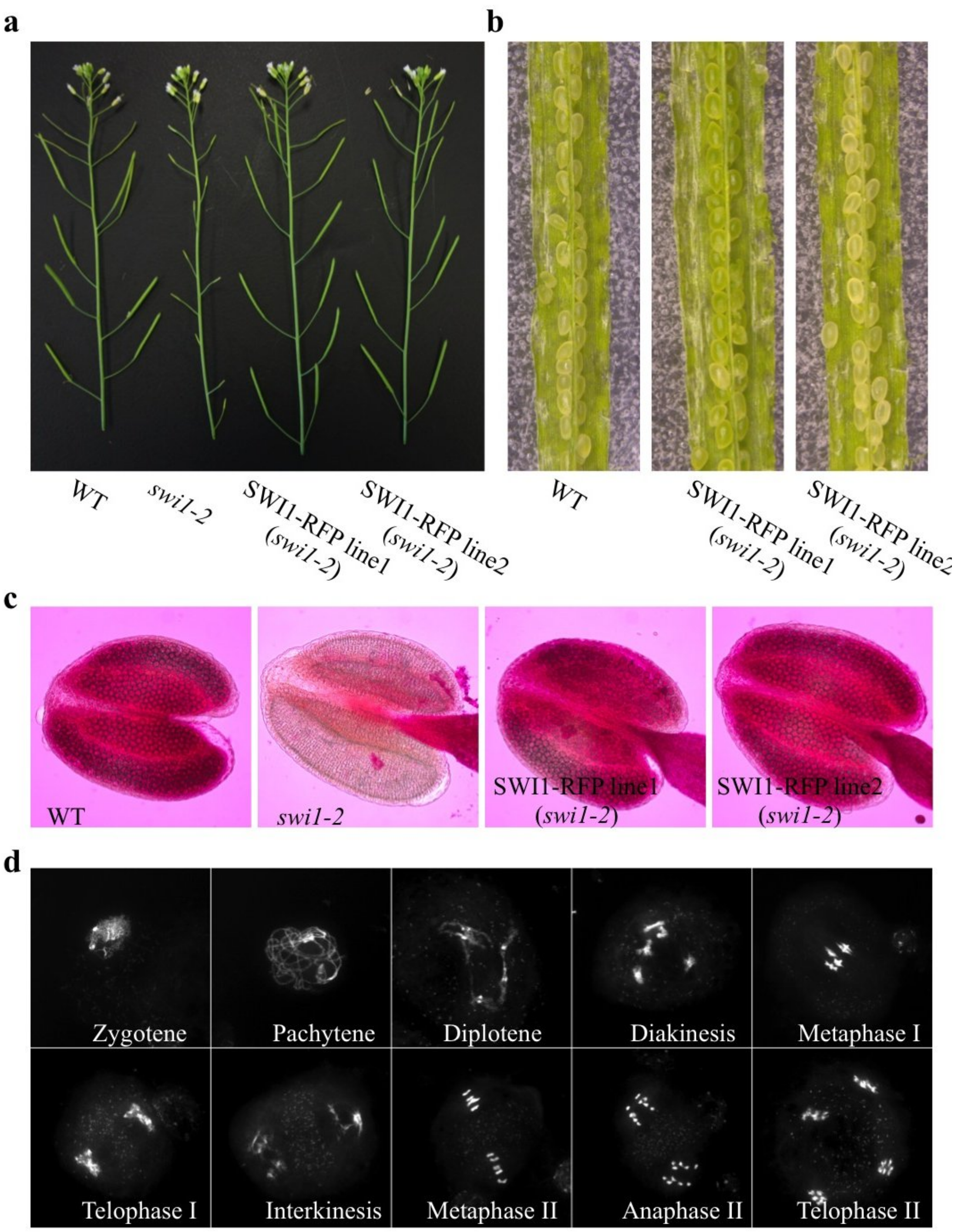


**Supplementary Figure 5**

**SWI1-RFP fully complements the meiotic defects in *swi1* mutants.** (a) Main branches of the wildtype (WT), *swi1-2* and two *SWI1-RFP* lines. (b) Seed sets in siliques of the wildtype (WT) and two *SWI1-RFP* lines. (c) Peterson staining of anthers for the wildtype (WT), *swi1-2* and *SWI1-GFP* lines. (d) Chromosome spread analysis of male meiocytes in *SWI1-RFP* line #1 (*swi1-2*) reveals a wild-type meiotic program.

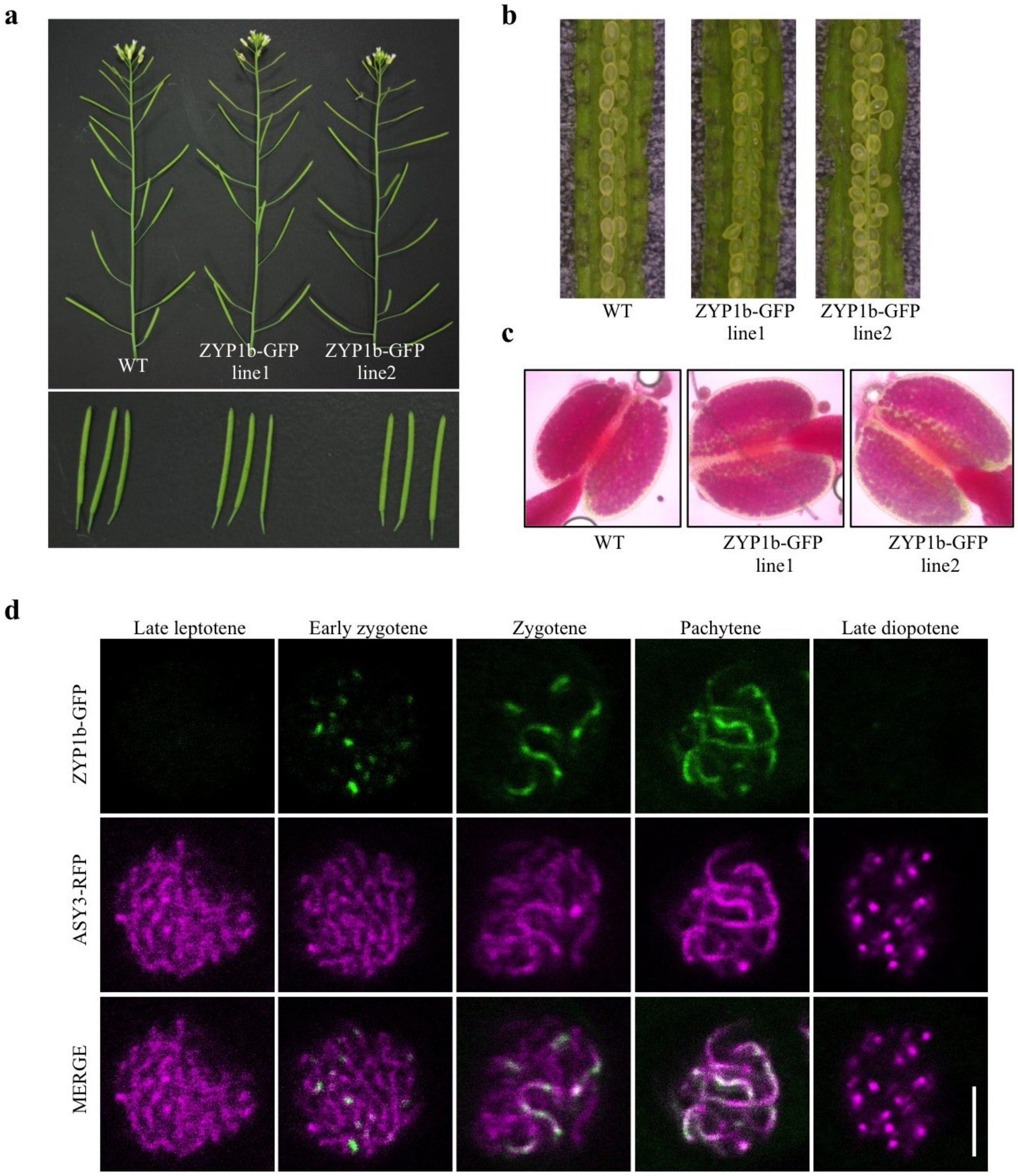


**Supplementary Figure 6**

**ZYP1b-GFP is a good reporter for staging and has no dominant effect on plants.**

(a) Main branches of the wildtype (WT) and two *ZYP1b-GFP* lines in wildtype bacground. (b) Seed sets in siliques of the WT and two *ZYP1b-GFP* lines. (c) Peterson staining of anthers for the WT and and two *ZYP1b-GFP* lines. (d) Co-localization of ZYP1b-GFP with ASY3-RFP in the male meiocytes of wildtype shows that ZYP1b-GFP specifically localizes to synaptic regions during prophase I. Bar: 5 μm.

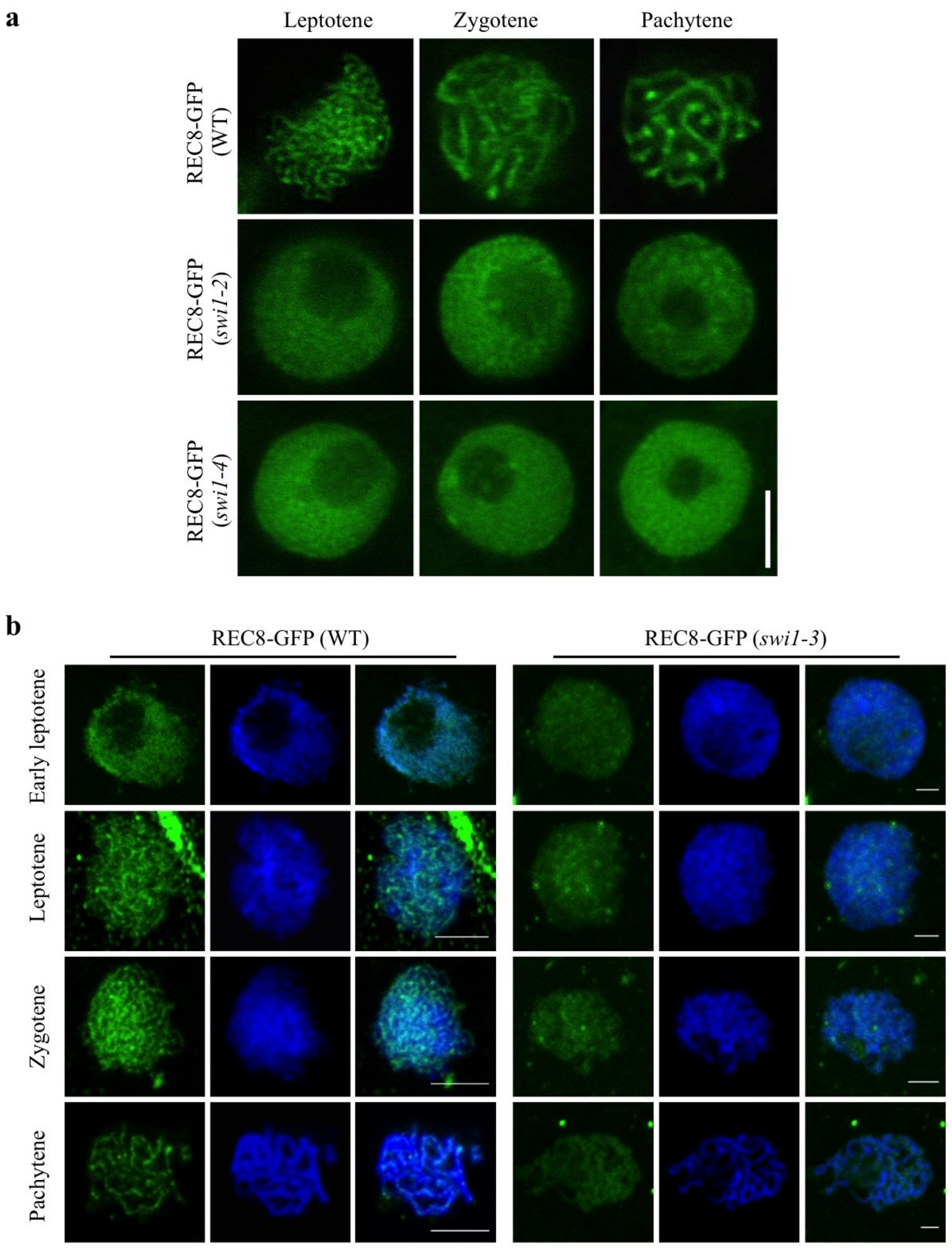


**Supplementary Figure 7**

**Cohesion establishment is compromised in different *swi1* alleles. (a)** Localization of REC8-GFP was analyzed by using laser confocal microscopy during prophase I of male meiocytes in the wildtype (WT), *swi1-2 and swi1-4*. Bar: 5 µm. (b) Immunolocalization of REC8-GFP in the male meiocytes of the wildtype and *swi1-3* mutants during prophase I. Anti-GFP antibody was used for detecting REC8-GFP.

Bar: 5 μm.

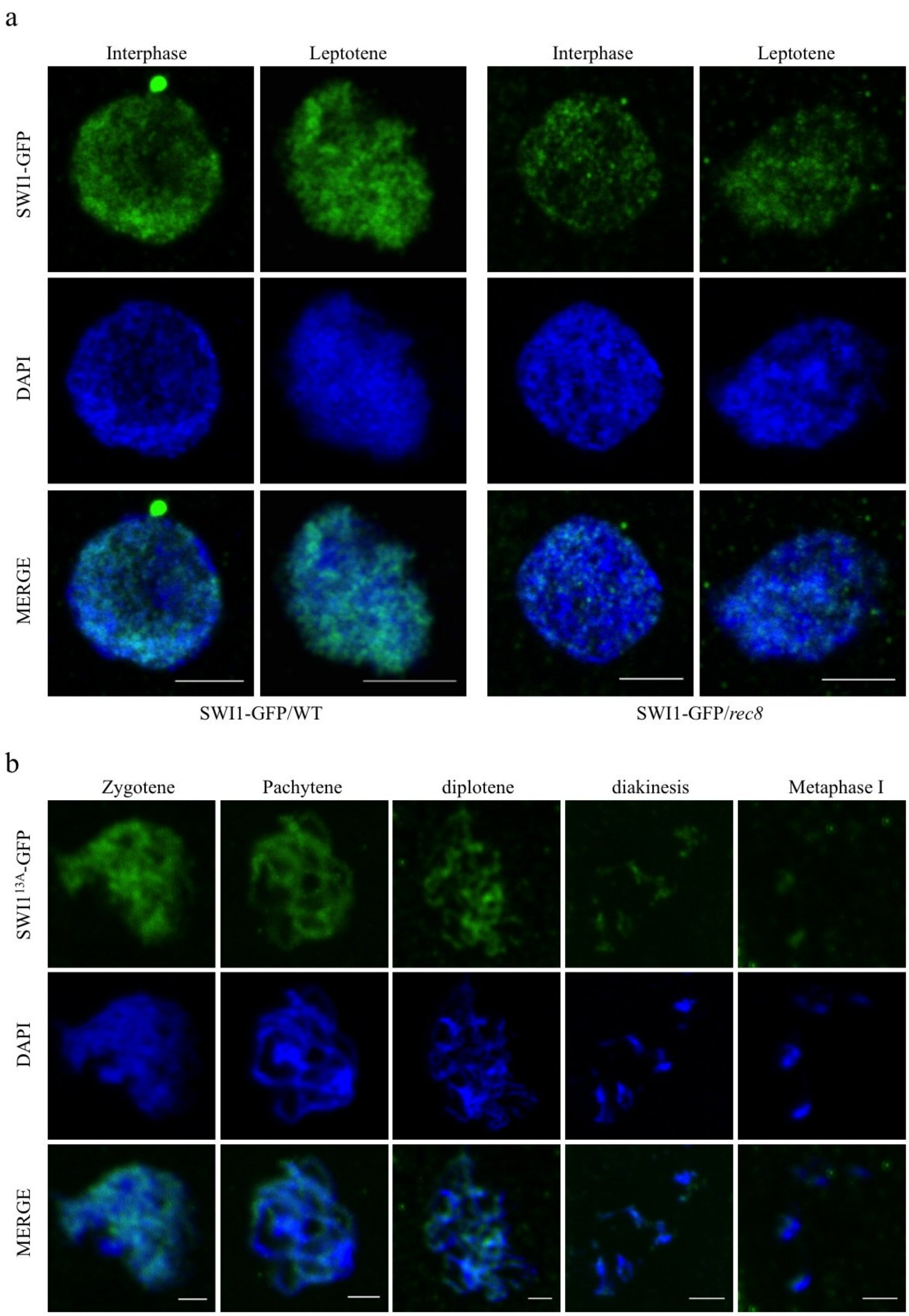


**Supplementary Figure 8**

**Immunolocalization of SWI1-GFP and SWI1$^{13A}$-GFP.** (a) Immunolocalization of SWI1-GFP in the wildtype (WT) and *rec8* mutants. Bar: 5 µm. (b)

Immunolocalization of SWI1$^{13A}$-GFP in the wildtype. Bar: 5 μm. Anti-GFP antibody was used for detecting SWI1.

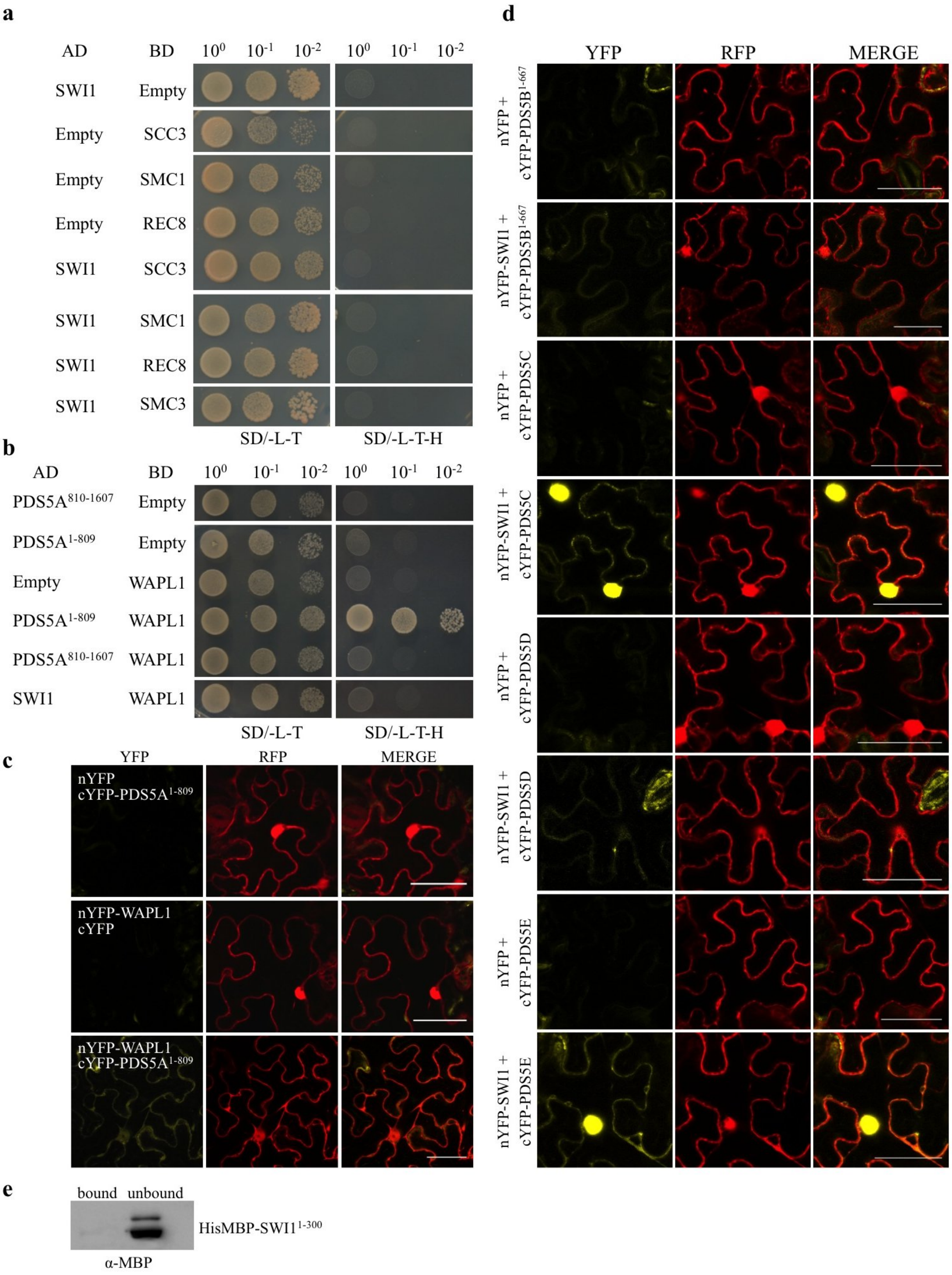


### Supplementary Figure 9

**Interaction analyses of SWI1, PDS5, and WAPL.** (a) Yeast two-hybrid interaction assay of SWI1 with the core cohesin subunits SMC1, SMC3, REC8 and SCC3. (b) Yeast two-hybrid analysis of the interactions of WAPL1 with PDS5 and SWI1. (c)

BiFC interaction assay of WAPL1 with PDS5A. Bar: 50 µm. (d) BiFC interaction assay of SWI1 with PDS5B, PDS5C, PDS5D, and PDS5E. (e) Immunoprecipitation for HisMBP-SWI1$^{1\text{-}300}$ only using GST binding beads showing no unspecific binding.

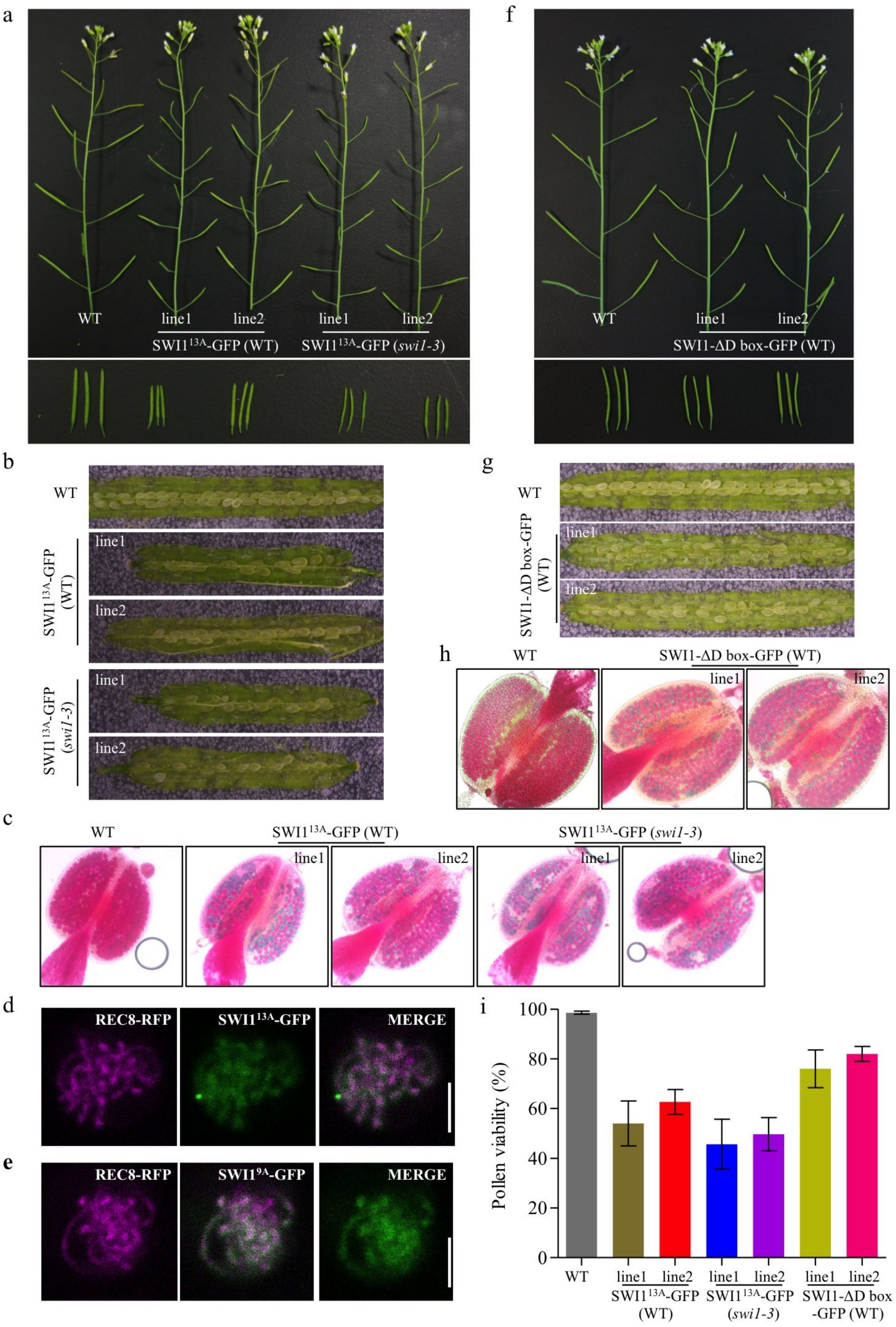


**Supplementary Figure 10**

**Fertility of plants harboring the SWI1$^{13A}$-GFP and SWI1-ΔD box-GFP constructs.** (a) Main branches (upper panel) and siliques (lower panel) of the wildtype (WT) and two *SWI1$^{13A}$-GFP* lines in both WT and *swi1-3* muatnt background. (b) Seed sets in siliques of the WT and two *SWI1$^{13A}$-GFP* lines in both WT and *swi1-3* muatnt background. (c) Peterson staining of anthers for the WT and and two *SWI1$^{13A}$-GFP* lines in both WT and *swi1-3* muatnt background. (d, e) Co-localization of REC8-RFP and SWI1$^{13A}$-GFP (d) and SWI1$^{9A}$-GFP (e) in the male meiocytes of *swi1-3* mutant at pachytene. Bar: 5 μm. (f) Main branches (upper panel) and siliques (lower panel) of the wildtype (WT) and two *SWI1-ΔD box-GFP* lines in WT background. (g) Seed sets in siliques of the WT and two *SWI1-ΔD box-GFP* lines in WT background. (h) Peterson staining of anthers for the WT and and two *SWI1-ΔD box-GFP* lines in WT background. (i) Quantification of the pollen viability for the plants shown in (a) and (f).

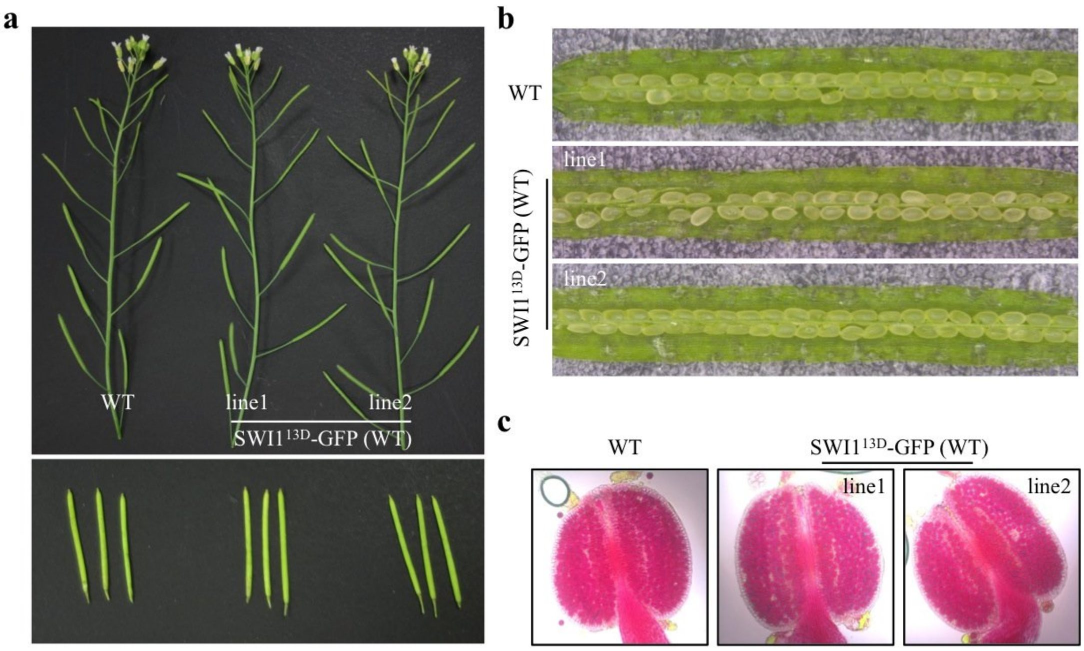


**Supplementary Figure 11**

**Fertility of plants harboring the SWI1$^{13D}$-GFP construct.** (a) Main branches (upper panel) and siliques (lower panel) of the wildtype (WT) and two *SWI1$^{13D}$-GFP* lines in wildtype (WT) background. (b) Seed sets in siliques of the WT and two *SWI1$^{13D}$-GFP* lines. (c) Peterson staining of anthers for the WT and and two *SWI1$^{13D}$-GFP* lines.

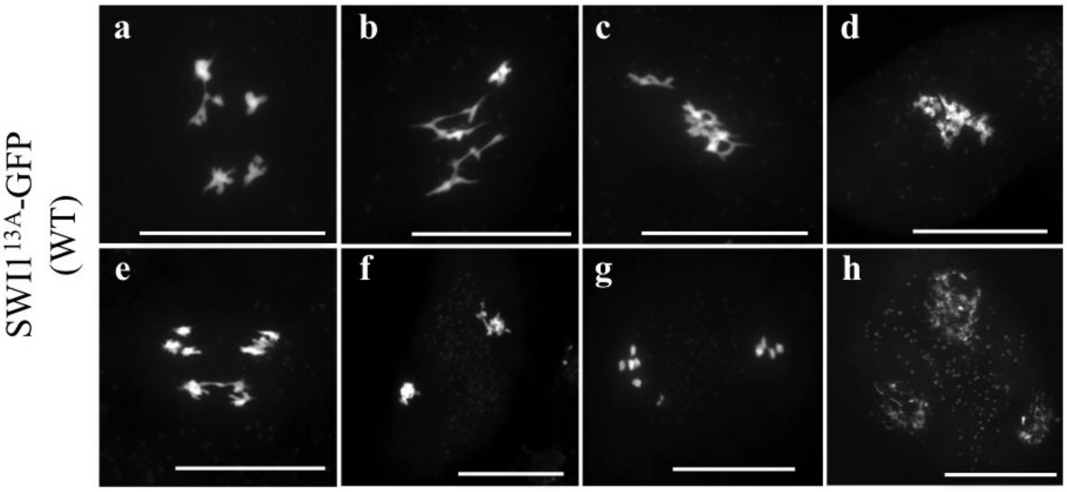


**Supplementary Figure 12**

**Chromosome spread analysis of male meiocytes in *SWI1$^{13A}$-GFP*/WT plants.** (a, b) diakinesis-like stage; (c, d) metaphase I-like stage; (e) anaphase I; (f, g) late telophase I or interkinesis; (f) Terad-like stage. Bar: 20 μm.

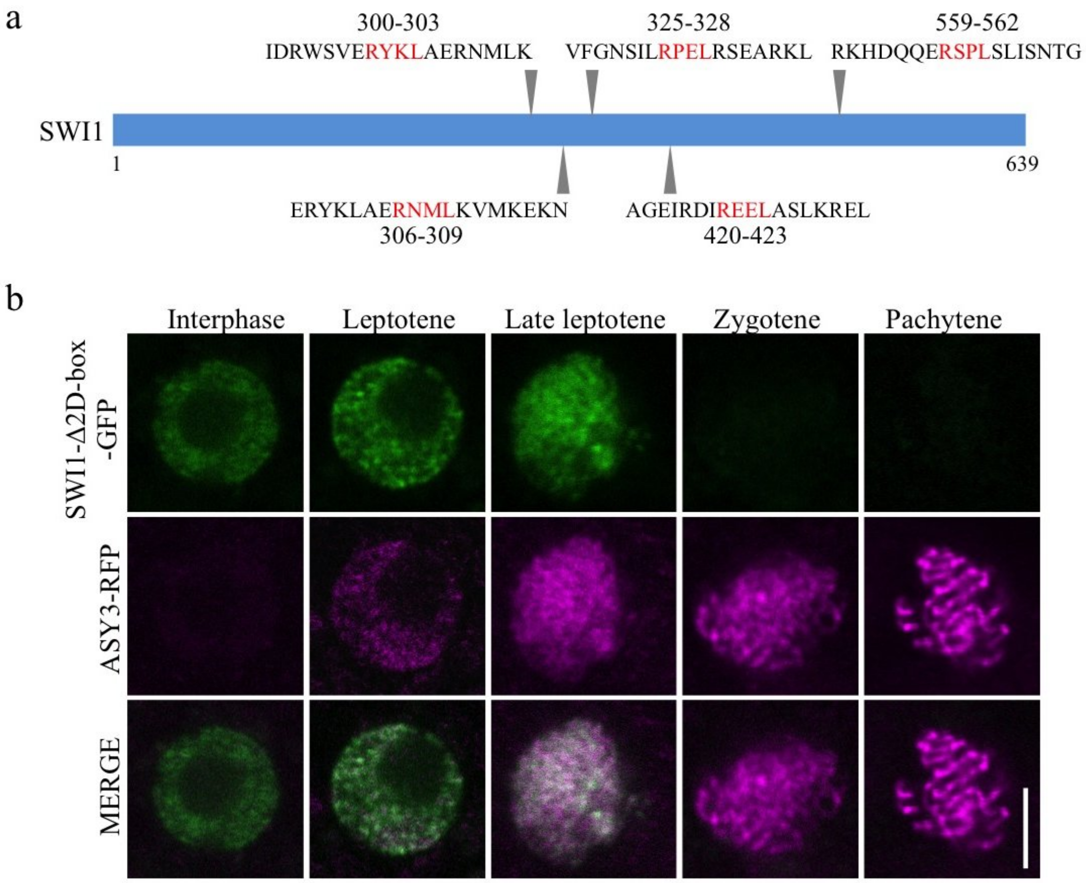


**Supplementary Figure 13**

**Mutation of two conserved destruction box (D-box) has no impact on the localization pattern of SWI1.** (a) Scheme of SWI1 coding region. Arrowheads indicate the positions of five putative D-boxes. (b) Localization analysis of SWI1-Δ2D-box-GFP (306-309 and 559-562) in the male meiocytes during prophase I. Bar: 5 μm.

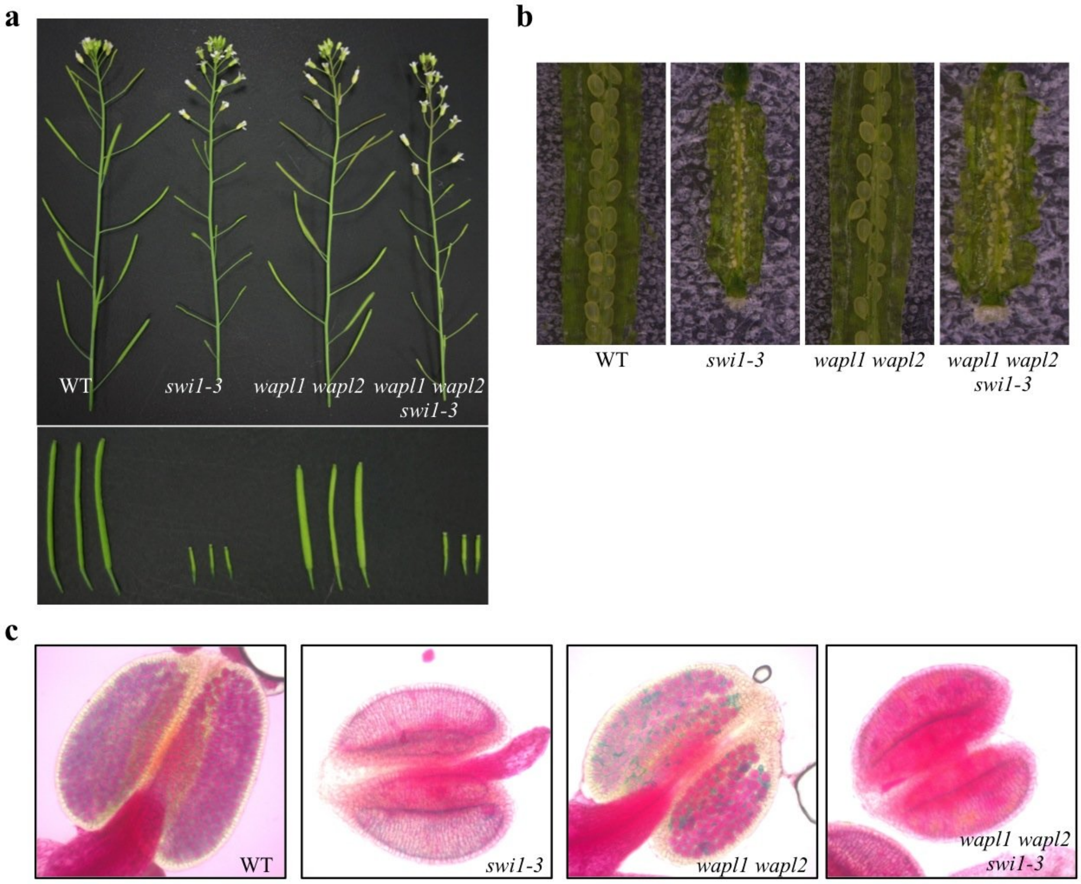


**Supplementary Figure 14**

**The absence of *WAPL1 WAPL2* does not restore the fertility of *swi1-3* mutants.**

(a) Main branches (upper panel) and siliques (lower panel) of the wildtype (WT), *swi1-3*, *wapl1 wapl2* and *wapl1 wapl2 swi1-3* mutants. (b) Seed sets in siliques of the wildtype (WT), *swi1-3*, *wapl1 wapl2* and *wapl1 wapl2 swi1-3* mutants. (h) Peterson staining of anthers for the wildtype (WT), *swi1-3*, *wapl1 wapl2* and *wapl1 wapl2 swi1-3* mutants.

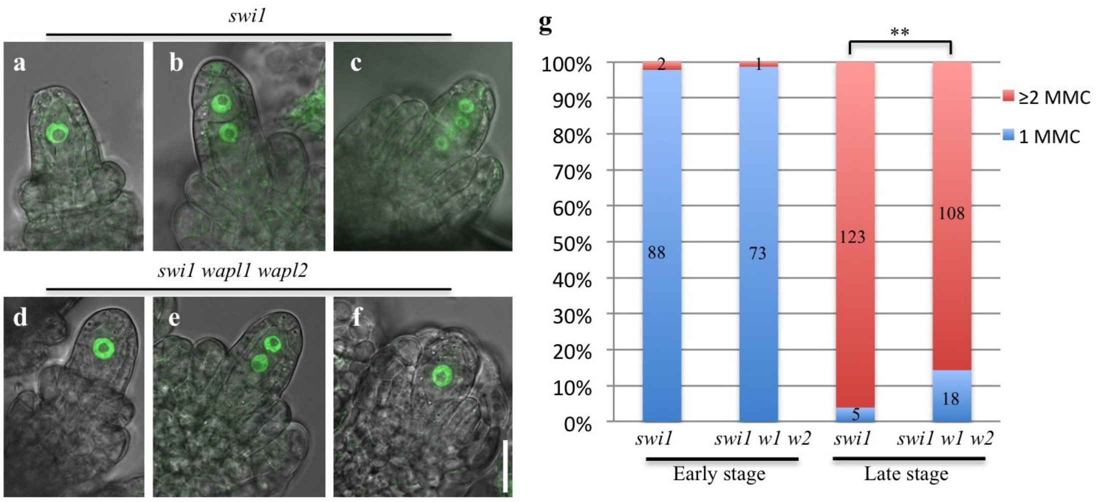


**Supplementary Figure 15**

**The formation of single megaspore mother cell (MMC) in *swi1* mutants is partially rescued by depletion of WAPL.** REC8-GFP was used as a maker for the counting of MMC. (a, d) Early premeiotic ovule with short integument primordia harboring one MMC. (b, e) Further developed ovules with elongated integuments containing two MMCs. (c) Older ovules than shown in (b) encompassing four MMCs. (f) Ovule at the similar stages as shown in (b, e) with only one MMC. Bar: 20 µm. (g) Statistical analysis of the number of MMCs at early (as in a, d) and late stages (as in b, c, e, f) in *swi1* and *swi1 wapl1 wapl2* (*swi1 w1 w2*) mutants. The numbers on the columns denote the amount of ovules counted. **, P< 0.01 (Chi-squared test).

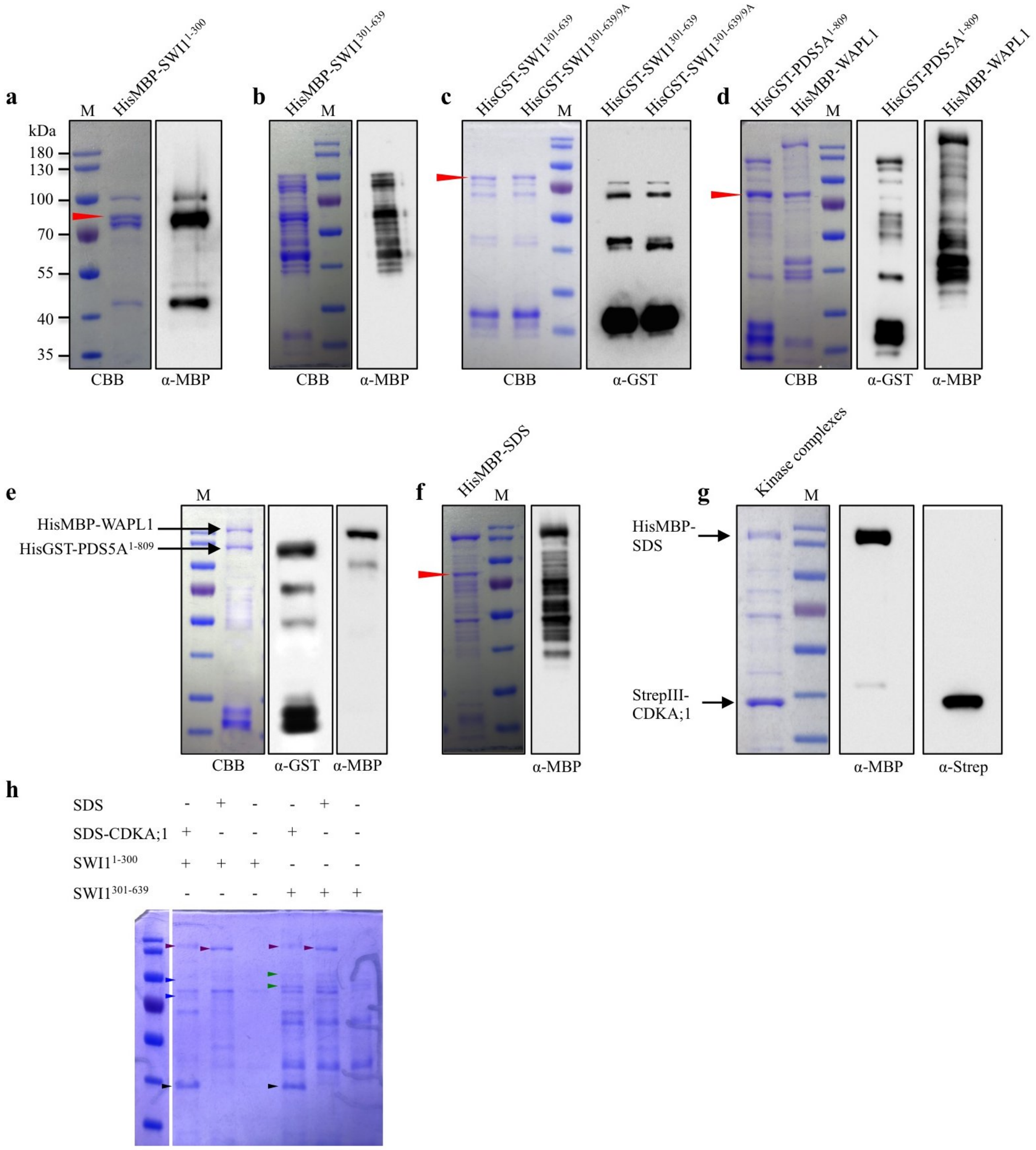


### Supplementary Figure 16

**CBB stained gels of all purified proteins from *Escherichia Coli* used in this research.** (a-c) CBB staining and western blot confirmation of purified HisMBP-SWI1$^{1\text{-}300}$ (a), HisMBP-SWI1$^{301\text{-}639}$ (b), HisGST-SWI1$^{301\text{-}639}$ and HisMBP-SWI1$^{301\text{-}639/9A}$ (c), HisMBP-PDS5A$^{1\text{-}809}$ and HisMBP-WAPL1 (d), HisMBP-PDS5A$^{1\text{-}809}$-HisMBP-WAPL1 heterodimers (e), HisMBP-SDS (f) and CDKA;1-SDS complexes (g) from *E.coli*. The arrowheads indicate the bands of unspecific protein binding generally to Ni-NTA beads. (h) CBB staining of the proteins after kinase reaction of SWI1 with CDKA;1-SDS complexes. The purple, blue, green or black arrowheads denote the main bands of SDS, $SWI1^{1-300}$, $SWI1^{301-639}$ or CDKA;1 proteins, respectively.

**Supplementary table 1.** Phosphorylated sites in SWI1.

| Experiments | Proteins | Positions | Amino acid | Localization probabilities | peptides and Phospho Probabilities |
|---|---|---|---|---|---|
| replicate 1 | SWI1 1-300; AT5G51330.1 | 22 | S | 0,99989 | ISS(1)PSSPTLNVAVAHIR |
| | SWI1 1-300; AT5G51330.1 | 166 | T | 0,998469 | REVVSQPAY(0.002)NT(0.998)R |
| | SWI1 301-639; AT5G51330.1 | 395 | T | 1,00000 | EAGVKDPYWT(1)PPPGWK |
| | SWI1 301-639; AT5G51330.1 | 447 | T | 0,89016 | KEEEELVIMT(0.11)T(0.89)PNSCVTSQNDNLMTPAK |
| | SWI1 301-639; AT5G51330.1 | 544 | S | 0,99992 | VVNKGNQITES(1)PQNR |
| | SWI1 301-639; AT5G51330.1 | 560 | S | 1,00000 | KHDQQERS(1)PLSLISNTGFR |
| | SWI1 301-639; AT5G51330.1 | 597 | S | 0.34000 | ICRPVGMFAWPQLPALAAATDT(0.037)NAS(0.478)S(0.34)PS(0.117)HR |
| | SWI1 301-639; AT5G51330.1 | 606 | S | 1,00000 | QAYPS(1)PFPVKPLAAK |
| replicate 2 | SWI1 1-300; AT5G51330.1 | 22 | S | 0,99998 | ISS(1)PSSPTLNVAVAHIR |
| | SWI1 1-300; AT5G51330.1 | 166 | T | 0,893899 | REVVS(0.001)QPAY(0.106)NT(0.894)R |
| | SWI1 301-639; AT5G51330.1 | 395 | T | 1,00000 | EAGVKDPYWT(1)PPPGWK |
| | SWI1 301-639; AT5G51330.1 | 447 | T | 0,60282 | KEEEELVIMT(0.354)T(0.603)PNS(0.035)CVTSQNDNLMTPAK |
| | SWI1 301-639; AT5G51330.1 | 544 | S | 0,99997 | VVNKGNQITES(1)PQNR |
| | SWI1 301-639; AT5G51330.1 | 560 | S | 0,98042 | KHDQQERS(0.98)PLS(0.02)LISNTGFR |
| | SWI1 301-639; AT5G51330.1 | 597 | S | 0,70145 | ICRPVGMFAWPQLPALAAATDTNAS(0.073)S(0.701)PS(0.225)HR |
| | SWI1 301-639; AT5G51330.1 | 606 | S | 0,99999 | QAYPS(1)PFPVKPLAAK |

Phosphorylated peptides of SWI1 were identified by mass spectrometry analysis after subjecting SWI1 to *in vitro* kinase assays with CDKA;1-SDS complexes, as shown in Fig. S9d. No phosphorylated peptides were found in reactions without CDKA;1. Results from two independent biological replicates are shown. Data are available via ProteomeXchange Consortium with the identifier PXD009959.

**Supplementary Table 2.** Primers used in this research.

| Purpose | Prmer name | sequence |
|---|---|---|
| SWI1 reporter | gSWI1-F | TTGACATTGTGAGAGTAACG |
| | gSWI1-R | AACTAGTCTAGAGAACGGGT |
| | gSWI1-attB1-F | GGGGACAAGTTTGTACAAAAAAGCAGGCTTAGCACTTTATGGTTTTTCCG |
| | gSWI1-attB2SmaI-R | GGGGACCACTTTGTACAAGAAAGCTGGGTTTCACCCGGGAACGTTGAAGAGATT CTTGG |
| ASY3 reporter | gASY3-F | TTTGAGAACTCCACTTTACTGCGT |
| | gASY3-R | CTGCTACTATCTTGTCGTCTTCTC |
| | gASY3-attB1-F | GGGGACAAGTTTGTACAAAAAAGCAGGCTTAAAAACATTACTTCCCCTACCAAA |
| | gASY3-attB2-SmaI-R | GGGGACCACTTTGTACAAGAAAGCTGGGTTTCACCCGGGATCATCCCTCAAACAT TCTGCGA |
| ZYP1b reporter | gZYP1b-F | GAAATCAGATGAGCCCTTCCTTAA |
| | gZYP1b-R | GGGAACTGACTTTGTGTGGTAGAC |
| | gZYP1b-attB1-F | GGGGACAAGTTTGTACAAAAAAGCAGGCTTAGAAATCAGATGAGCCCTTCC |
| | gZYP1b-attB2-R+T | GGGGACCACTTTGTACAAGAAAGCTGGGTTTTATCAATCAAATGCATAGGGATC |
| | gZYP1b-AscI-F | AAGCATGGCGCGCCGGTAATAAGAGAAGCGAGCA |
| | gZYP1b-AscI-R | AAGCATGGCGCGCCACCAAGACGAGATTCTTTCA |
| | GFP-AscI-F | AAGCATGGCGCGCCAGGTGGCGGTGGATCAGGCGG |
| | GFP-AscI-R | AAGCATGGCGCGCCCAGACCCTCCACCTCCCTTGT |
| Y2H | SMC1-EcoRI-F | CCGGAATTCATGCCTGCGATACAATCCCCATCG |
| | SMC1-SalI-R | ACGCGTCGACTCACGATTCTTGGTAGTTCCTAAGG |
| Y2H | SMC3-NcoI-F | CATGCCATGGGAATGTTTATCAAGCAGGTTATAATCG |
| | SMC3-BamHI-R | CGCGGATCCTCAGGTATCGTGGGACTGATCTTTC |
| Y2H | REC8-EcoRI-F | CCGGAATTCATGTTGAGACTGGAGAGTTTGATAG |
| | REC8-SalI-R | ACGCGTCGACTTACATGTTGGGTCCTCTTGCAATG |
| Y2H | SCC3-EcoRI-F | CCGGAATTCATGGAAGACAGTCCTCAAGGCCTTA |
| | SCC3-SalI-R | ACGCGTCGACTCAGTGTCCCTTGGACCGTTCACCC |
| Y2H | SWI1-EcoRI-F | CCGGAATTCATGAGTAGTACGATGTTCGTGAAAC |
| | SWI1-XhoI-R | CCGCTCGAGTCAAACGTTGAAGAGATTCTTGG |
| Y2H and protein expression | SWI1-attB1-F | GGGGACAAGTTTGTACAAAAAAGCAGGCTTCATGAGTAGTACGATGTTCGTGAA A |
| | SWI1-300aa-attB2-R | GGGGACCACTTTGTACAAGAAAGCTGGGTTTCACCTCTCAACAGACCATCTATCA |
| | SWI1-301aa-attB1-F | GGGGACAAGTTTGTACAAAAAAGCAGGCTTCTACAAACTAGCTGAGAGGAACAT G |
| | SWI1-639aa-attB2-R | GGGGACCACTTTGTACAAGAAAGCTGGGTTTCAAACGTTGAAGAGATTCTTGGG |
| Y2H and protein expression | AtPDS5A 1-809aa-F | ATGGCTCAGAAGCCGGAGGAACAGTTGAAAG |
| | AtPDS5A 1-809aa-R | CTACTTAACCAACGTCTTGATCCCATATATCTTC |
| | AtPDS5A 810-1607aa-F | CTGAAGATATATGGGATCAAGACGTTGGTT |
| | AtPDS5A 810-1607aa-R | CTATATTGCTGTCCTCGAGATTGACTTACCCAC |
| | AtPDS5A 1-809-attB1 F | GGGGACAAGTTTGTACAAAAAAGCAGGCTTCATGGCTCAGAAGCCGGAGGAACA G |
| | AtPDS5A 1-809-attB2 R | GGGGACCACTTTGTACAAGAAAGCTGGGTTCTACTTAACCAACGTCTTGATCCCA |
| | AtPDS5A 810-1607-attB1 F | GGGGACAAGTTTGTACAAAAAAGCAGGCTTCCTGAAGATATATGGGATCAAGAC |
| | AtPDS5A 810-1607-attB2 R | GGGGACCACTTTGTACAAGAAAGCTGGGTTCTATATTGCTGTCCTCGAGATTGAC |
| Y2H and protein expression | WAPL1-CDS-F | ATGATAATTGTAAAACTAACGGCCAATCGC |
| | WAPL1-CDS-R | CTACGGTGATTTGCAGGATTCAATCACTCCC |
| | WAPL1-CDS-attB1-F | GGGGACAAGTTTGTACAAAAAAGCAGGCTTCATGATAATTGTAAAACTAACGGC C |
| | WAPL1-CDS-attB2-R | GGGGACCACTTTGTACAAGAAAGCTGGGTTCTACGGTGATTTGCAGGATTCAATC |
| Y2H | OsAM1-CDS F | ATGGACGCGGAGATGGCGGCTCCTGCGCTTG |

| | | |
|---|---|---|
| Y2H | OsAM1-CDS R | TCAGCAGTAGGACGGAGTGGCCAGTGCCAGCTC |
| | OsAM1-attB1-F | GGGGACAAGTTTGTACAAAAAAGCAGGCTTCATGGACGCGGAGATGGCGGCTCC |
| | OsAM1-attB2-R | GGGGACCACTTTGTACAAGAAAGCTGGGTTTCAGCAGTAGGACGGAGTGGCCAG |
| | ZmAM1-CDS F | ATGGACGTAGAGACGGTGCAGGCGGGTCCTG |
| | ZmAM1-CDS R | TCAGCAGTAGGATGGAGTAGCCAGGGCCAGCTC |
| | ZmAM1-attB1-F | GGGGACAAGTTTGTACAAAAAAGCAGGCTTCATGGACGTAGAGACGGTGCAGGC |
| | ZmAM1-attB2-R | GGGGACCACTTTGTACAAGAAAGCTGGGTTTCAGCAGTAGGATGGAGTAGCCAG |
| Dephospho mutagenesis | SWI1 S22/25A-F | GCTCCGTCGGCTCCGACTTTGAATGgtaaactactga |
| | SWI1 S22/25-R | AGAGATTTTCCCGGCGGTGGTTTCT |
| | SWI1 S52A-F | GCTCCGGAAAATCTTAAATCGATTAGAG |
| | SWI1 S52-R | TCTCTGAGGAAGAATCGAAGCATCG |
| | SWI1 S173A-F | GCTCCGGAGGGAAAGTGCTCGTCTGAG |
| | SWI1 S173-R | AGCAGCGCGACAGAGACGAGTATTG |
| | SWI1 T242A-F | GCTAAACAAGAGGCAAAGGAGATAACTA |
| | SWI1 T242-R | GCCTCCTATTTCATTCCCATCATCA |
| | SWI1 S261A-F | GCTAGTACTGAGAGACTCGCTCAGAAAG |
| | SWI1 S261-R | TTCAATCAGCTTTCTCTTACGATT |
| | SWI1 T395A-F | GCTCCTCCACCTGGTTGGAAGCTTGGTG |
| | SWI1 T395-R | CCAGTAAGGATCTTTAACTCCTGCT |
| | SWI1 T447A-F | GCTCCTAATTCTTGTGTTACTAGTCAG |
| | SWI1 T447-R | AGTCATGATAACAAGCTCCTCCTCT |
| | SWI1 T461A-F | GCTCCAGCAAAGgtaagagctcgaaaca |
| | SWI1 T461-R | CATCAGATTATCATTCTGACTAGTA |
| | SWI1 T515A-F | GCTCCTTTGCTACTAGAGGATTCACCAC |
| | SWI1 T515-R | CTCTGTTGAGTCTGGCTTTTTAGGA |
| | SWI1 S522A-F | GCTCCACCAATACAGACACTAGAAGGAG |
| | SWI1 S522-R | ATCCTCTAGTAGCAAAGGTGTCTCT |
| | SWI1 S544A-F | GCTCCTCAAAACAGAGAAAAAGGAAGGA |
| | SWI1 S544-R | CTCTGTGATTTGGTTACCCTTGTTC |
| | SWI1 S560A-F | GCTCCACTTTCACTAATAAGCAACACTG |
| | SWI1 S560-R | TCTTTCTTGTTGATCATGCTTCCTT |
| | SWI1 S597A-F | GCTCCAAGTCACAGACAAGCCTACCCAT |
| | SWI1 S597-R | AGAAGCATTAGTATCAGTAGCAGCA |
| | SWI1 S606A-F | GCTCCTTTTCCAGTCAAGCCACTTGCAG |
| | SWI1 S606-R | TGGGTAGGCTTGTCTGTGACTTGGC |
| genotyping for *swi1-2* | SWI1-CAPS-F | AACAAGAGGCAAAGGAGATAAC |
| | SWI1-CAPS-R | TTTTCAGCAGATCAGCCGTAGA |
| genotyping for *swi1-3* | SAIL_654_C06 LP | ACTCATCACCGCTTGATTCTG |
| | SAIL_654_C06 RP | TGATACTGCACACGCAATCTC |
| genotyping for *swi1-4* | GABI_206H06 LP | CTCCCAGATTCATTAAATGCG |
| | GABI_206H06 RP | CTAGAAACCCAGAAACCCCAG |
| Phosphomimic mutagenesis | SWI1 S22/25D-F | GACCCGTCGGACCCGACTTTGAATGgtaaactactga |
| | SWI1 S22/25-R | AGAGATTTTCCCGGCGGTGGTTTCT |
| | SWI1 S52D-F | GACCCGGAAAATCTTAAATCGATTAGAG |
| | SWI1 S52-R | TCTCTGAGGAAGAATCGAAGCATCG |
| | SWI1 S173D-F | GACCCGGAGGGAAAGTGCTCGTCTGAG |
| | SWI1 S173-R | AGCAGCGCGACAGAGACGAGTATTGTAC |
| | SWI1 T395D-F | GACCCTCCACCTGGTTGGAAGCTTGGTG |

| | | |
|---|---|---|
| Phosphomimic mutagenesis | SWI1 T395-R | CCAGTAAGGATCTTTAACTCCTGCT |
| | SWI1 T447D-F | GACCCTAATTCTTGTGTTACTAGTCAG |
| | SWI1 T447-R | AGTCATGATAACAAGCTCCTCCTCT |
| | SWI1 T461D-F | GACCCAGCAAAGgtaagagctcgaaaca |
| | SWI1 T461-R | CATCAGATTATCATTCTGACTAGTA |
| | SWI1 T515D-F | GACCCTTTGCTACTAGAGGATTCACCAC |
| | SWI1 T515-R | CTCTGTTGAGTCTGGCTTTTTAGGA |
| | SWI1 S544D-F | GACCCTCAAAACAGAGAAAAAGGAAGGA |
| | SWI1 S544-R | CTCTGTGATTTGGTTACCCTTGTTC |
| | SWI1 S560D-F | GACCCACTTTCACTAATAAGCAACACTG |
| | SWI1 S560-R | TCTTTCTTGTTGATCATGCTTCCTT |
| | SWI1 S597D-F | GACCCAAGTCACAGACAAGCCTACCCAT |
| | SWI1 S597-R | AGAAGCATTAGTATCAGTAGCAGCA |
| | SWI1 S522D-F | GACCCACCAATACAGACACTAGAAGGAG |
| | SWI1 S522-R | ATCCTCTAGTAGCAAAGGTGTCTCT |
| | SWI1 S606D-F | GACCCTTTTCCAGTCAAGCCACTTGCAG |
| | SWI1 S606-R | TGGGTAGGCTTGTCTGTGACTTGGC |
| BiFC constructs | AtPDS5A-attB1 F1 | GGGGACAAGTTTGTACAAAAAAGCAGGCTTCATGGCTCAGAAGCCGGAGGAACAG |
| | AtPDS5A-attB4 R1-T 809aa | GGGGACAACTTTGTATAGAAAAGTTGGGTGCTTAACCAACGTCTTGATCCCA |
| | AtPDS5B-CDSattB1-F1 | GGGGACAAGTTTGTACAAAAAAGCAGGCTTCATGGAGAAAACTCCGACGCAG |
| | AtPDS5B-CDSattB4-R1_T | GGGGACAACTTTGTATAGAAAAGTTGGGTGGCCACAAAGCTGATTCAAAAG |
| | AtPDS5C-CDSattB1-F | GGGGACAAGTTTGTACAAAAAAGCAGGCTTCATGTCGGATTCTGATAAAGAG |
| | AtPDS5C-CDSattB4-R_T | GGGGACAACTTTGTATAGAAAAGTTGGGTGTCGCTTCCTCTTCTTACCGG |
| | AtPDS5D-CDSattB1-F | GGGGACAAGTTTGTACAAAAAAGCAGGCTTCATGCAAAGTGCCCTAATTCCATC |
| | AtPDS5D-CDSattB4-R_T | GGGGACAACTTTGTATAGAAAAGTTGGGTGTGACTTTCTCTTCTCTTCATC |
| | AtPDS5E-CDSattB1-F | GGGGACAAGTTTGTACAAAAAAGCAGGCTTCATGGGTCCTCTTGTCGAAGCG |
| | AtPDS5E-CDSattB4-R_T | GGGGACAACTTTGTATAGAAAAGTTGGGTGTGGATCAACCTCAAGCACC |
| | AtSWI1-CDS-attB3-F | GGGGACAACTTTGTATAATAAAGTTGTAATGAGTAGTACGATGTTCGTGAAA |
| | AtSWI1-639-attB2-R-T | GGGGACCACTTTGTACAAGAAAGCTGGGTTAACGTTGAAGAGATTCTTGGG |
| | AtSWI1-300-attB2-R-T | GGGGACCACTTTGTACAAGAAAGCTGGGTTCCTCTCAACAGACCATCTATCA |
| | WAPL1-attB3-F | GGGGACAACTTTGTATAATAAAGTTGTAATGATAATTGTAAAACTAACGGCC |
| | WAPL1-attB2-R-T | GGGGACCACTTTGTACAAGAAAGCTGGGTTCGGTGATTTGCAGGATTCAATC |

**Supplementary Video 1**

**Dynamics of REC8-GFP in wild-type plants.** Live cell imaging of REC8-GFP was performed in male meiocytes of wild-type plants. Video starts at leptotene stage and runs for 25 h with scan intervals of 30 mins. Bar: 10 µm.

**Supplementary Video 2**

**Dynamics of REC8-GFP in *wapl1 wapl2* mutants.** Live cell imaging of REC8-GFP was performed in male meiocytes of *wapl1 wapl2* mutants. Video starts at leptotene stage and runs for 25 h with scan intervals of 30 mins. Bar: 10 µm.

**Supplementary Video 3**

**Dynamics of REC8-GFP in *swi1* mutants.** Live cell imaging of REC8-GFP was performed in male meiocytes of *swi1* mutants. Video starts at early zygotene-like stage and runs for 21 h with scan intervals of 15 mins. Bar: 10 µm.

**Supplementary Video 4**

**Dynamics of REC8-GFP in *swi1 wapl1 wapl2* mutants.** Live cell imaging of REC8-GFP was performed in male meiocytes of *swi1 wapl1 wapl2* mutants. Video starts at early zygotene-like stage and runs for 21 h with scan intervals of 15 mins. Bar: 10 µm.

**Supplementary Video 5**

**Dynamics of REC8-GFP in wild-type plants.** Live cell imaging of REC8-GFP was performed in male meiocytes of wild-type plants. Video starts at early leptotene stage and runs for 30 h with scan intervals of 15 mins. Bar: 10 µm.

**Supplementary Video 6**

**Dynamics of REC8-GFP in *SWI1*$^{13A}$*-GFP*/WT plants.** Live cell imaging of REC8-GFP was performed in male meiocytes of *SWI1*$^{13A}$*-GFP*/WT plants. Video starts at early leptotene stage and runs for 35 h with scan intervals of 15 mins. Bar: 10 µm.
